# Genetic correlation between sea age at maturity and iteroparity in Atlantic salmon

**DOI:** 10.1101/412288

**Authors:** Tutku Aykanat, Mikhail Ozerov, Juha-Pekka Vähä, Panu Orell, Eero Niemelä, Jaakko Erkinaro, Craig R. Primmer

## Abstract

Genetic correlations in life history traits may result in unpredictable evolutionary trajectories if not accounted for in life-history models. Iteroparity (the reproductive strategy of reproducing more than once) in Atlantic salmon (*Salmo salar*) is a fitness trait with substantial variation within and among populations. In the Teno River in northern Europe, iteroparous individuals constitute an important component of many populations and have experienced a sharp increase in abundance in the last 20 years, partly overlapping with a general decrease in age structure. The physiological basis of iteroparity bears similarities to that of age at first maturity, another life history trait with substantial fitness effects in salmon. Sea age at maturity in Atlantic salmon is controlled by a major locus around the *vgll3* gene, and we used this opportunity demonstrate that the two traits are genetically correlated around this genome region. The odds ratio of survival until second reproduction was up to 2.4 (1.8-3.5 90% CI) times higher for fish with the early-maturing *vgll3* genotype (*EE*) compared to fish with the late-maturing genotype *(LL*). The association had a dominance architecture, although the dominant allele was reversed in the late-maturing group compared to younger groups that stayed only one year at sea before maturation. *Post hoc* analysis indicated that iteroparous fish with the *EE* genotype had accelerated growth prior to first reproduction compared to first-time spawners, across all age groups, while this effect was not detected in fish with the *LL* genotype. These results broaden the functional link around the *vgll3* genome region and help us understand constraints in the evolution of life history variation in salmon. Our results further highlight the need to account for genetic correlations between fitness traits when predicting demographic changes in changing environments.

## Introduction

Since being formally described, multivariate evolution has been well incorporated into quantitative genetic frameworks, covering predictions under diverse theoretical scenarios (Lande, 1979; Wagner, 1989; Houle, 1991; Roff, 1996; Griswold & Whitlock, 2003; Chevin *et al.*, 2010; Wang *et al.*, 2010). Accordingly, the theory suggests genetic correlation between traits can constrain the pace and efficacy of natural selection (Lande & Arnold, 1983; Roff, 1996; Orr, 2000). When correlated characters have contrasting fitness trajectories in the adaptive landscape, “the climb” towards the local fitness peak will be restricted, and this will result in suboptimal fitness of populations (Lande, 1982). The principles of the multivariate theory of evolution have been successfully applied to many fields, such as animal and plant breeding, multi-trait artificial selection (Kadarmideen *et al.*, 2003; Careau *et al.*, 2010; Chen *et al.*, 2011; Kause *et al.*, 2011; Weigel *et al.*, 2017), and epidemiology (Lee *et al.*, 2012; Sanchez-Guillen *et al.*, 2012; Bulik-Sullivan *et al.*, 2015; Gratten & Visscher, 2016; Schnurr *et al.*, 2016; Hammerschlag *et al.*, 2017). Measuring multi-trait evolution in wild populations also has substantial importance. The magnitude and sign of genetic correlation between fitness traits may facilitate or constrain adaptive evolution by conflicting trait co-evolution, or so-called trade-offs (Roff, 1996; Sheldon *et al.*, 2003; Hellmann & Pineda-Krch, 2007; Agrawal & Stinchcombe, 2009; Duputie *et al.*, 2012; Chirgwin *et al.*, 2015). A handful of studies have investigated the genetic relationships between fitness-related traits in the wild (Sheldon *et al.*, 2003; Theriault *et al.*, 2007; Carlson & Seamons, 2008; Nussey *et al.*, 2008; Robinson *et al.*, 2009; Clements *et al.*, 2011; Lane *et al.*, 2011; Santure *et al.*, 2013), most of which were confined to well-studied populations in isolated or historically monitored settings with a scope to demonstrate the evolution of antagonistic genetic correlation between fitness traits (Roff, 1996). Indeed, accurately estimating genetic relatedness in the wild is challenging and requires either well-established pedigrees (Kruuk *et al.*, 2000) or the genotyping of large numbers of genetic markers in large datasets (Ritland, 1996; Yang *et al.*, 2010; Robinson *et al.*, 2013; Berenos *et al.*, 2014). Thus, examples from wild populations are limited. Furthermore, multivariate genetic models are much more data-demanding than univariate models, making estimating genetic correlations more difficult than measuring univariate additive genetic variation (Roff, 1996; Nussey *et al.*, 2008; Wilson *et al.*, 2010).

Despite these challenges, measuring multivariate trait evolution in the wild has substantial implications for conservation and management efforts, e.g., to better predict population responses to changing environmental conditions (Etterson & Shaw, 2001; Hellmann & Pineda-Krch, 2007; Chirgwin *et al.*, 2015). In general, if a significant predictor of population demographic structure is left unaccounted, predictions may be inaccurate (Walters & Maguire, 1996; Dunlop *et al.*, 2009). In this sense, any potential genetic correlation between fitness-related traits will mediate the evolutionary response of populations and alter fitness optima (hence absolute fitness), perhaps towards an unexpected direction, and therefore should not be overlooked. Also important but often overlooked in wild populations is the fact that genetic correlations between fitness traits may uncover the functional and physiological basis of the correlation at the molecular level (Stearns *et al.*, 1991; Storz *et al.*, 2015). This can help illuminate the response of populations to ecosystem level processes and facilitate understanding how ecological dynamics shape trait evolution (Storz *et al.*, 2015).

Sea age at first maturity and iteroparity are two life history traits with profound effects on both reproductive output and survival (Fleming, 1996; Fleming & Einum, 2010) in Atlantic salmon (*Salmo salar*) and other salmonid fish species (Christie *et al.*, 2018). The two traits are phenotypically correlated, such that repeat spawning is more prevalent in fish maturing at a younger age, likely as a result of less energy investment in the first reproductive event and hence better post-reproduction recovery (Jonsson *et al.*, 1991b; 1997; Niemelä *et al.*, 2006a; Penney & Moffitt, 2013). We therefore hypothesized that the traits were also genetically correlated. We employed a gene-trait association approach to test for the existence of a genetic correlation between sea age at first maturity and iteroparity in Atlantic salmon. More specifically, we took advantage of a recently reported large-effect locus controlling age at first maturity in the region of the *vgll3* gene on Atlantic salmon chromosome 25 (Ayllon *et al.*, 2015; Barson *et al.*, 2015) and explored whether the same region explained variation in iteroparity.

## Materials and Methods

### Study site and life history of repeat spawning salmon in the Teno River

Located in far-north Europe (68–70°N, 25–27°E), the Teno River runs between Finland and Norway and drains north into the Barents Sea at the Tana Fjord (Fig. 1). The river supports one of the world’s largest wild Atlantic salmon populations and accounts for up to 20% of the riverine Atlantic salmon catches in Europe (ICES 2013). The Teno River supports over 20 Atlantic salmon sub-populations (Vähä *et al.*, 2017) with notably high genetic and life-history variation within and between populations (Vähä *et al.*, 2007; Aykanat *et al.*, 2015; Vähä *et al.*, 2017; Erkinaro *et al.*, 2018). Age at smoultification (i.e., number of years spent in fresh water prior to outward migration to the sea) varies between two and eight years, while the time spent in the marine environment prior to first maturation, called sea age at maturity, varies from one to five years. In addition, a proportion of individuals have an iteroparous life history, in which individuals attempt to reproduce more than once, and sometimes up to three times, in their adult life (Niemelä *et al.*, 2006a; Erkinaro *et al.*, 2018) (see also Figure S1).

**Figure 1:**
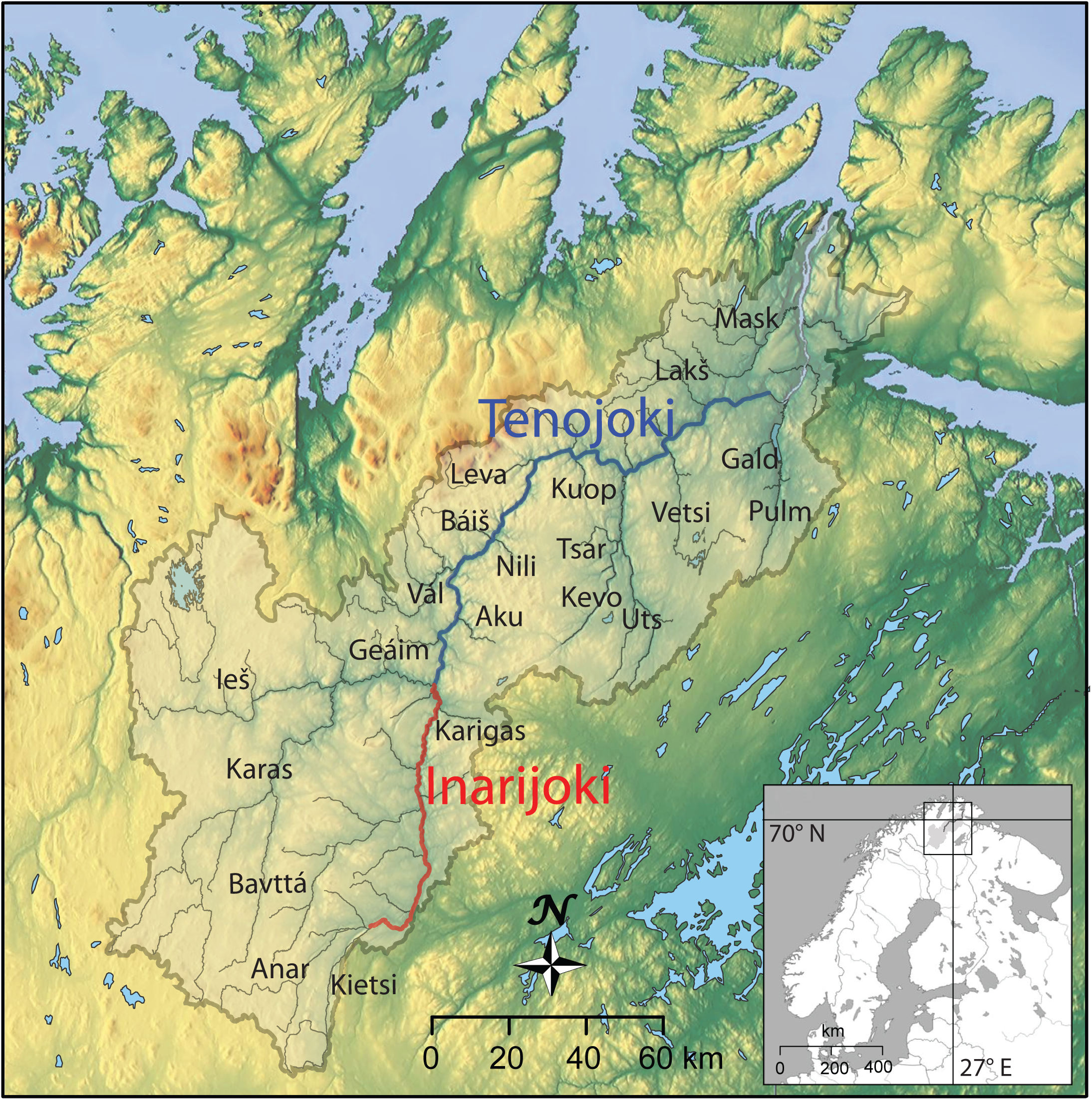
Map of the study system. River sections with blue and red colours indicate Tenojoki and Inarijoki, respectively. (See Table S2 for full river names.)

In the Teno River, individuals who survive their first spawning event most often spend a full year at sea for conditioning (i.e., alternate year repeat spawner) before returning to the river in the following summer (Figure S1). Three major repeat-spawning strategies comprise more than 85% of the repeat spawner life history types in the mainstem Teno River: individuals who spend one, two or three winters at sea before first spawning, then spend an additional full year at sea re-conditioning following the first spawning event, and then return to the spawning grounds the next summer as a repeat spawner (Erkinaro *et al.*, 2018). These three repeat spawning strategies were the focus of our analyses and are referred to as 1S1, 2S1 and 3S1, respectively.

The relative proportion of a specific previous spawner life history is mostly correlated to the abundance of the respective maiden year cohort and to the fact that one full year of re-conditioning before returning to fresh water to spawn again is the most common post-spawning life history in the high latitude Teno River (Niemelä *et al.*, 2006a; Erkinaro *et al.*, 2018). In the Teno River, the average proportion of repeat-spawner salmon, based on catch statistics, is 5% over the last four decades, but the abundance and proportion have been increasing in every major repeat-spawner life history type since the early 2000s, which cannot be explained only by the increase in the abundance of the respective maiden cohort (Figure S2, Erkinaro *et al.,* 2018).

### Sample collection

Atlantic salmon scales can be used to infer a number of life-history characteristics, including smolt age (years spent in fresh water from hatch sea migration), sea age to first spawning event (years spent at sea prior to first spawning, also referred to as age at maturity), and iteroparity (evidence of, and time between, multiple spawning migrations) (Erkinaro et al. 2018; Figure S1). The fish samples used in this study were from a scale collection archive of Atlantic salmon from the Teno River system, which spans more than four decades and is maintained by Natural Resources Institute Finland (LUKE; former Finnish Game and Fisheries Research Institute). Scale samples were composed of riverine catches of anadromous adult Atlantic salmon. These scales have been collected by co-operating, trained fishers of the Teno River system, who have also been recording phenotypic traits (e.g., fish total length, sex) and the location, date and method of capture of the fish. As defined by standard guidelines, scales were sampled from below the adipose fin, just above the lateral line, where the age of the fish can be most reliably inferred (ICES, 2011). The collected scales were later dried and archived in paper envelopes at the Teno River Fisheries Research station of LUKE in Utsjoki, Finland. Trained technicians inferred the freshwater and sea age of fish using standard methods (ICES, 2011).

For this study, first-time spawners were defined as adults captured in fresh water whilst returning from the sea for their first reproduction attempt. In total, 643 first-time spawners representing three sea age at first maturity classes were included in this study (*N*_*1SW*_=228, *N*_*2SW*_=170, and *N*_*3SW*_=245, where 1*SW,* 2*SW* and 3*SW* denote the years (sea-winters) spent at sea before the first breeding attempt). The phenotypic and genotypic information of these individuals has previously been reported in (Aykanat *et al.*, 2015). The samples represented individuals captured between 2001 and 2003, along an ~130-km stretch of the Teno River mainstem, reaching ca. 210 km from the sea between 2001 and 2003 (see Aykanat *et al.* (2015) for details). Sampled fish were captured in the last four weeks of the fishing season, in August (two to four weeks after most individuals have entered the river), to minimize the number of fish from tributary and headwater populations (Erkinaro *et al.*, 2010). Using the abovementioned dataset, Aykanat *et al.* (2015) identified two sub-populations that have been subsequently identified to represent the Teno mainstem (Tenojoki, referred to as sub-population 1 and Inarijoki (sub-population 2) sub-populations (Pritchard *et al.*, 2018).

We further studied scales from 492 repeat spawner individuals (*N*_*1S1*_=225, *N*_*2S1*_=155, and *N*_*3S1*_=112) that had good DNA quality. Fish were selected non-randomly with respect to sea age and sex to balance samples from low proportion life history types. Most fish were captured between 2001 and 2008, but a small proportion of sampled fish were captured during later years (<8% sampled between 2008 and 2014. Table S1). Repeat spawner sampling spanned a broader time window within the fishing season than first-time spawners due to the much lower total number of repeat-spawner fish late in the season, since repeat spawners tend to return to breeding grounds earlier than the maiden fish ((Niemelä *et al.*, 2006b), Table S1).

### DNA extraction, sex determination and SNP genotyping by targeted sequencing

DNA extraction and sex determination of first-time spawners were carried out as described elsewhere (Johnston *et al.*, 2014; Aykanat *et al.*, 2015). For the repeat spawner group, DNA was extracted from one to two scales per individual using a QIAamp 96 DNA QIAcube HT Kit (Qiagen), following the manufacturer’s protocol, and with an initial proteinase K digestion step. Quality and concentration of all DNA extractions was assessed using a Nanodrop ND-1000 spectrophotometer (Thermo Fisher Scientific Inc.).

Repeat spawner samples were genotyped by targeted sequencing that included 194 SNP loci and the sex determination locus (*sdy*) as outlined in Aykanat *et al.* (2016) with minor modifications. Briefly, genomic regions were first amplified in two multiplex PCR reactions using site-specific primers with adapter sequences. After an SPRI bead clean-up to reduce short, non-specific reads, the PCR products of each individual were combined and re-amplified with adapter-specific primers containing Ion Torrent and sample-specific sequences. The PCR product was again purified by SPRI bead clean-up, quantified with Qubit 2.0 fluorimeter, and pooled in equimolar concentrations into one library (maximum 288 samples together). The pooled library was diluted for template preparation using the Ion PGM Hi-Q OT2 kit for Ion AmpliSeq DNA Library and OT2 for 200 bp reads and enrichment steps (ES) according to the manufacturer’s instructions. Finally, samples were sequenced using an Ion PGM Hi-Q sequencing kit and Ion 318 Chip 2 following the manufacturer’s guidelines.

SNPs in the targeted sequencing panel (N = 194) were described in Aykanat *et al.* (2016). The panel consists of putatively neutral and highly diverged SNPs between the Inarijoki and Tenojoki sub-populations (neutral module: N=136, outlier module: N=53) and potentially functionally important SNPs that are associated with sea age at maturity on chromosomes 9 and 25 (sea age module, N=5). The baseline and outlier modules allow for the quantification of population genetic parameters and reliably assign population of origin (Aykanat *et al.* (2016). The sea age module consists of four SNPs on chromosome 25, located in the genome region associated with age at maturity (Ayllon *et al.*, 2015; Barson *et al.*, 2015), of which two are missense SNPs in *vgll3* (*vgll3*_*Met54Thr*_ and *vgll3*_*Asn323Lys*_), one is a missense SNP in *akap11* (*akap11*_*Val214Met*_), and one is the SNP with the most significant association with age at maturity (i.e., *vgll3*_*TOP*_ from Barson *et al.,* 2015). In addition, the module has an SNP from the chromosome 9 region that exhibits a strong association with sea age at maturity prior to population structure correction (Barson *et al.*, 2015). This SNP is 34.5 kb away from and in complete linkage disequilibrium with the *SIX6*_*TOP*_ SNP from Barson *et al.* (2015) (termed *SIX6*_*TOP.LD*_ here). The majority of the SNP data for first-time spawners were taken from previous studies (Johnston *et al.*, 2014; Aykanat *et al.*, 2015; Barson *et al.*, 2015), with the exception of two missense SNPs, *vgll3*_*Met54Thr*_ and *vgll3*_*Asn323Lys*_, which were sequenced using the Ion Torrent platform as described above.

Unless otherwise noted, all statistical analyses were performed using R software v.3.2.5 (R Core Team 2013). Raw *fastq* output from the Ion Torrent server was scored using custom R scripts as outlined in Aykanat *et al.* (2016). Briefly, forward and reverse barcodes were used as identifiers to assign reads to individuals, followed by assigning within-individual data to each locus by matching reads to locus-specific primers (allowing for one mismatch). Reads were further refined by a sequence pattern match, this time to a 9-bp region surrounding each SNP locus, without a mismatch allowed. Finally, loci with coverage less than 13 were excluded, and the remainder were assigned a genotype as in Campbell *et al.* (2015). After excluding three SNPs with low genotyping rates (<75%), genotyping success on average was 96.5%. Molecular sexing was conducted by estimating read counts of the *sdy* gene (normalized to mean read count for every individual) using an arbitrary threshold as outlined in Aykanat *et al.* (2016).

### Genetic assignment of individuals using genetic baseline data

All sampled fish were captured as late in the season as possible in order to maximize the proportion of fish from focal populations (i.e., Tenojoki and Inarijoki). Due to the lower number of iteroparous individuals in the population, repeat-spawner individuals captured earlier in the season were included, which may have increased the proportion of fish from non-focal, tributary populations amongst this group (Table S1). Therefore, all samples underwent population assignment to enable exclusion of individuals potentially originating from non-focal populations using 175 non-sea-age associated SNPs that were successfully genotyped both in the dataset and in baseline samples. The baseline consisted of 23 sub-populations within the Teno system (Fig. 1). We first estimated the allele frequency distributions of the SNPs across the baseline populations and then estimated the likelihood of each individual to originate from each of those populations using a frequency-based method (see Table S2 for more details). The robustness of assignments was shown to be high using simulations, whereby random generation of 1000 genotypes per population showed high true assignment rates to Tenojoki and Inarijoki (93.2% and 91.2%, respectively); a negligible proportion of individuals from other populations were incorrectly assigned to these focal populations with high confidence (0.3% and 0.8%, respectively).

Out of 643 first-time spawners, 335 (52.1%) and 167 (26.0%) were assigned to Tenojoki and Inarijoki, respectively (78.1% in total), while 16.6% were unassigned, and 5.3% were assigned to other populations (Table S2). For 478 repeat-spawning individuals, 115 and 141 fish were assigned to Tenojoki and Inarijoki populations, respectively (53.6%, in total), leaving 137 fish (28.7%) unassigned and 85 fish (17.8%) confidently assigned to other populations in the Teno system (Table S2). The high proportion of misassigned fish in the repeat spawning category was expected, given that this group also included individuals sampled earlier in the fishing season (May-July). Individuals from these months included a higher proportion of fish captured in their non-natal spawning grounds. In contrast, the proportion of fish assigned to non-focal populations was similar between first-time spawners and repeat spawners in August (exact-test, p>0.05, Table S3). Following this assignment procedure, the final data set included 502 first-time spawners and 256 repeat spawners (Table S2).

### Testing genetic associations to iteroparity

In Atlantic salmon, sea age at first maturation (i.e., sea age) is strongly controlled by one genomic region (i.e., around the *vgll3* gene, approximately 28.65 Mb on chromosome 25, Barson *et al.*, 2015). An additional region, around the *six6* gene on chromosome 9, also exhibits a strong association with sea age at the population level, though the signal diminishes after correcting for structure (Barson *et al.*, 2015). Here, we tested if variation at five SNPs linked to sea age at first maturity on chromosome 9 and 25 (*vgll3*_*TOP*_, *vgll3*_*Met54Thr*_, *vgll3*_*Asn323Lys*_, *akap11*_*Val214Met,*_ *SIX6*_*TOP*_ (see above) were also linked to the repeat spawning life history. To do this, we first identified the most parsimonious null model that fit the data without genetic effects and then employed the genetic model over it. This allowed for us to avoid any biased inference that may have emerged as a result of unequal proportions of previous spawners within sea age groups, sex, populations or their higher-order interactions. We employed a general linear model with a binomial error structure, where repeat-spawner fish were coded as a Boolean variable “1”, first-time spawners were coded as “0”, and the allelic effect was an independent variable. Origin of population, sea age at first maturity, and sex were included in the model as cofactors. Using a semi-automatic model-scanning approach in the MuMIn package (Barton, 2018) in R (R Core Team 2013), a full-null model (all cofactors with all possible interactions, but without the genetic term) and reduced null models were compared by the corrected Akaike information criterion score (*AICc*), which is an *AIC* score with a stronger penalty for complex models. The null model explaining the data the best (with the lowest *AICc* score) was then used as the null hypothesis when testing the genetic effect. The genetic effect was then included in the optimum null model as an independent additive effect. If the alternative hypothesis was accepted over the null, we further refined the best genetic model(s) with a similar approach to that above, i.e., by testing models with all possible interactions with the genetic term over the null model. We also employed the genetic term as categorical model to test if any non-additive models explained the data better than the additive model.

Finally, to understand possible growth variation among genotypes associated with repeat spawning, length at capture was investigated as a function of genotype within each life history group (i.e., repeat and first-time spawners). We again employed a model evaluation approach as above, where a full-interaction model, consisting of sea age at first maturity, sex, and population, and reduced models were evaluated to find the most parsimonious model. The response variable, total length at capture, was log-normalized.

## Results

The best null model explaining the repeat-spawning life-history strategy (without genetic effects as a term) included population, sea age at first spawning, sex and the interaction between sex and population as factors (Table S4). The highest improvement in the fit of the model for explaining repeat spawning compared to the null model was obtained when SNPs in the chromosome 25 region associated with sea age at maturity were included in the model (Fig. 2). In this region, *akap11*_*Val214Met*_ showed the strongest association with the repeat spawner phenotype, followed by missense SNPs on *vgll3* gene (*vgll3*_*Met54Thr*_ and *vgll3*_*Asn323Lys*_ see Barson et al 2015) and *vgll3*_*TOP*_ loci (all *p* values<0.001, see Fig. 2). *SIX*_*TOP.LD*_ on chromosome 9 did not exhibit an association different from the other SNPs in the panel (Fig. 2).

**Figure 2:**
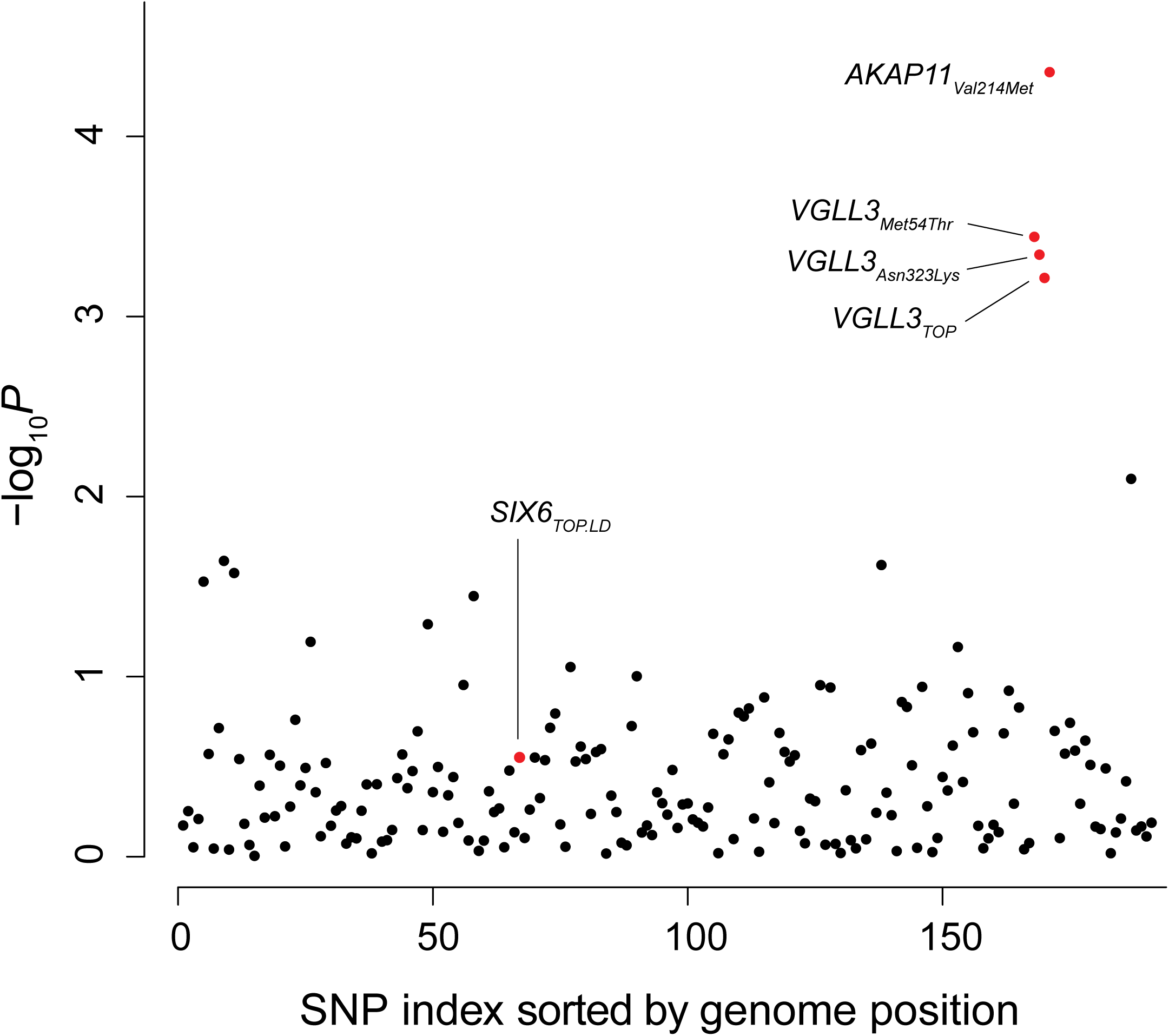
Goodness of the fit of the genetic models in 188 SNPs. P values indicate the significance of the fit of the genetic model relative to the null model. SNPs previously associated with age at maturity are labelled and coloured red.

In order to better understand the fine-scale genetic architecture of the association, we then explored genetic models with alternative parametrization of the genetic effect and evaluated the goodness of fit for each of the four SNPs in the chromosome 25 region. The missense polymorphism at *vgll3*_*Met54Thr*_ exhibited the highest fit across all loci (Table S5). In the best-supported model, the genetic architecture was consistent with non-additivity (SNP was coded as a categorical effect), which was dependent on sex as well as sea age at maturity (Table 1, Table S5, Table S6). Even though *vgll3*_*Met54Thr*_ explained the data better than other loci in the chromosome 25 region, the difference was not substantial. For example, when the dataset was resampled 1000 times (random sampling within sex and sea age groups with replacement), models including *vgll3*_*Met54Thr*_ were the best-fitting models in 42% of cases, while models with *vgll3*_*TOP*_, *vgll3*_*Asn323Lys*_, and *akap11*_*Val214Met*_ were better fitting 21.9%, 14.9% and 21.2% of the time, respectively. Therefore, we cannot clearly distinguish between the importance of the SNPs in the region with respect to their association with repeat spawner prevalence.

**Table 1:** Coefficients of the most parsimonious model for surviving to second spawning (repeat spawning) in Atlantic salmon at the *vgll3*_Met54Thr_ locus. Sea age denotes the sea age at first maturity. E and L denote alleles linked to the early and late age at maturity, respectively. See Table S5 for a full list of all plausible models.

| Coefficient | Estimate | Std.Error | z-value | p-value |
| --- | --- | --- | --- | --- |
| Inarijoki | 0.656 | 0.452 | 1.45 | 0.147 |
| Tenojoki | 0.682 | 0.523 | 1.30 | 0.192 |
| Sex (Male) | -0.008 | 0.243 | -0.03 | 0.973 |
| Sea age | -0.686 | 0.194 | -3.53 | 0.000 |
| SNP (EL) | -0.786 | 0.508 | -1.55 | 0.122 |
| SNP (EE) | 0.050 | 0.621 | 0.08 | 0.936 |
| Sea Age : SNP (EL) | 0.706 | 0.228 | 3.10 | 0.002 |
| Sea Age : SNP (EE) | 0.323 | 0.347 | 0.93 | 0.351 |
| Tenojoki : Sex (Male) | -0.837 | 0.346 | -2.42 | 0.015 |

The odds ratio of surviving to second spawning was estimated as a function of *vgll3*_*Met54Thr*_ genetic variation using the most parsimonious model. Generally, the genotype associated with earlier first maturity at sea (*EE*, *E* denotes the allele associated with early first-time maturation) was more likely to survive to second spawning than the genotype associated with later first-time maturation (*LL*, *L* denotes the allele associated with late first-time maturation) (Fig. 3, Table 1). This effect was age-dependent with a slight but not significant effect of population and a sex × population interaction (Fig. 3, Table 1, Table S7). The odds ratio of surviving to second spawning differed less between the genotypes of 1S1 fish: the *EE* genotype was 1.19 (0.91-1.91, 90% CI, *p*=0.14) and 1.24 (1.00-1.57, 90% CI, *p*=0.051) times more likely to survive to second spawning compared to the *LL* and *EL* genotypes, respectively. Older sea age groups exhibited stronger differences between genotypes, where the odds ratio between *LL* and *EE* genotype fish was 1.50 (1.12-2.05, 90% CI, *p*=0.012) and 2.03 (1.07-3.57, 90% CI, *p*=0.035) for 2S1 and 3S1 fish, respectively (Fig. 3, Table S7). The genetic architecture was consistent with dominance, which, however, appears to be age-dependent (Fig. 3, Table S7). In 3S1 fish, the genetic architecture significantly deviated from additivity (*p*=0.014, Table S7), where the *L* allele was dominant, while in 1S1 fish, the genetic architecture marginally deviated from additivity in the reverse direction (*p*=0.053).

**Figure 3:**
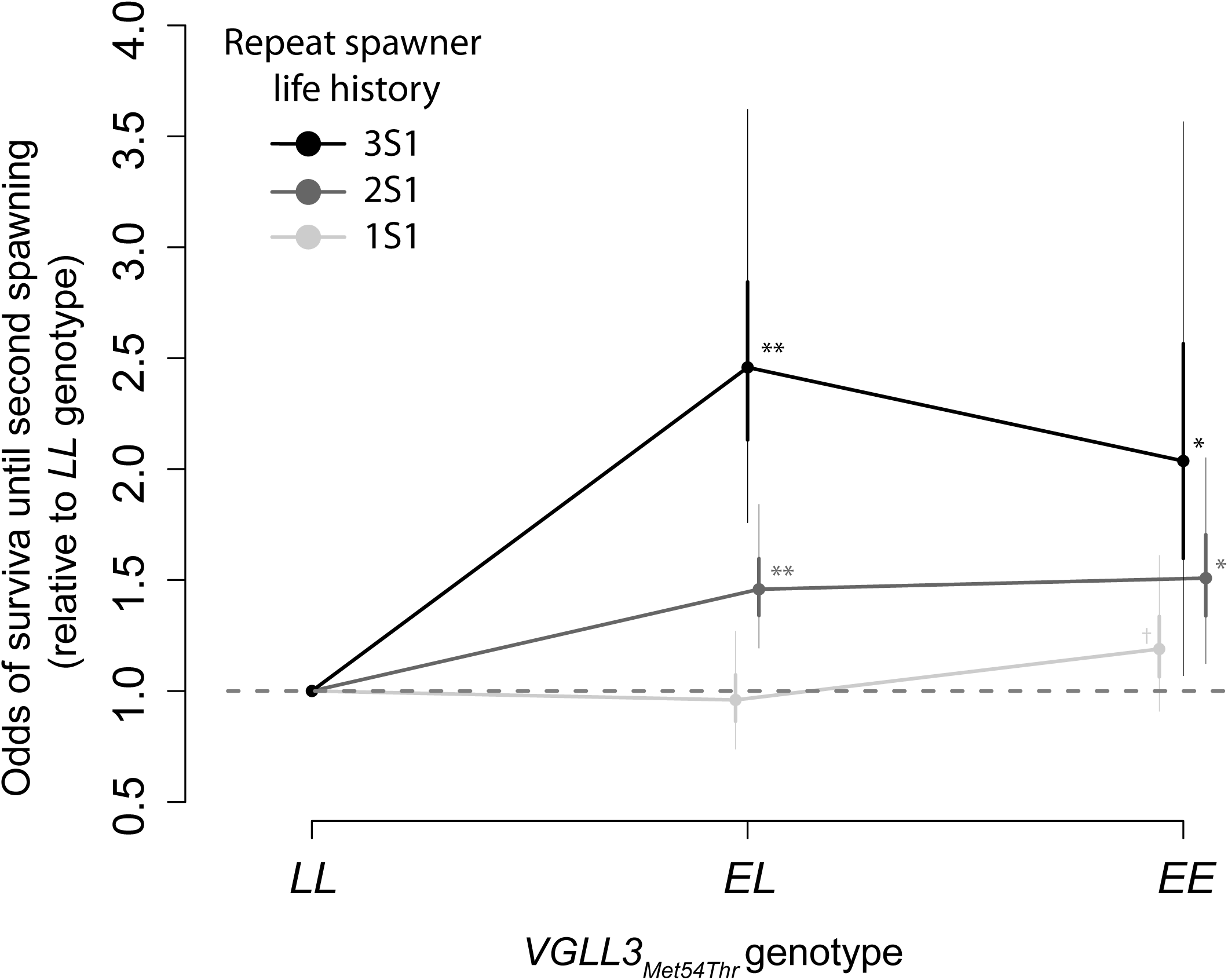
Odds of surviving to second spawning (repeat spawning) in *EE* and *EL* genotypes, relative to *LL* genotype, at the *vgll3*_Met54Thr_ locus, as estimated by 10000 parametric permutations. Dots, thick lines, and thin lines denote median estimates, 50% CIs, and 90% CIs, respectively. Asterisks denote significance, as calculated by the proportion of permutations whose odds of survival were greater than that of the *LL* genotype (* p<0.05, ** p<0.001; † denotes the proportion of permutations whose odds of survival were greater than that of the EL genotype at p=0.051). (See Table 1 for full model specifications.)

Finally, we tested for the allelic effect of *vgll3*_*Met54Thr*_ on total length at capture separately for the first-time spawner and repeat spawner groups. In the first-time spawners, the allelic effect was parametrized as in the most parsimonious model without any higher-order interactions, and the *L* allele was positively associated with length (Table 2, Table S8). This relationship was similar to that reported by Barson *et al.* (2015) for the *vgll3*_*TOP*_ locus. In contrast, there was no allelic association with length in the repeat spawner group (Table 2, Table S8). These two results combined suggest that allele-specific growth differences were offset in the repeat-spawner fish.

**Table 2:** Coefficients of the most parsimonious model explaining variation in total length at capture in first-time spawner and repeat spawner groups. The first-time spawner model includes *vgll3*_Met54Thr_, in which alleles are coded additively (i.e., *EE*=1, *EL*=2, and *LL* = 3). Length is log-normalized. See Table S8 for a list of all plausible models.

|  | Coefficient | Estimate | Std. Error | *t*-value | *p*-value |
| --- | --- | --- | --- | --- | --- |
| First-time<br>spawners | Intercept | 3.755 | 0.022 | 171.80 | <0.001 |
|  | SNP | 0.021 | 0.005 | 3.90 | <0.001 |
|  | Population | 0.223 | 0.044 | 5.09 | <0.001 |
|  | Sea age | 0.258 | 0.011 | 22.83 | <0.001 |
|  | Sex | -0.031 | 0.032 | -0.98 | 0.329 |
|  | Population : Sea age | -0.072 | 0.018 | -4.06 | <0.001 |
|  | Population : Sex | -0.185 | 0.052 | -3.56 | <0.001 |
|  | Sea age : Sex | 0.053 | 0.022 | 2.39 | 0.017 |
|  | Population : Sea age : Sex | 0.056 | 0.027 | 2.10 | 0.037 |
| Repeat<br>spawners | Intercept | 4.226 | 0.016 | 267.37 | <0.001 |
|  | Population | 0.080 | 0.011 | 7.19 | <0.001 |
|  | Sea age | 0.139 | 0.008 | 16.41 | <0.001 |
|  | Sex | 0.095 | 0.023 | 4.18 | <0.001 |
|  | Sea age : Sex | -0.049 | 0.012 | -4.00 | <0.001 |

## Discussion

Surviving following the first breeding event to potentially reproduce additional times (i.e., repeat spawning) is an important life history feature of Atlantic salmon. The trait is closely connected to fitness. Hence, understanding the ecological dynamics around the trait variation and its genetic underpinnings is important for predicting Atlantic salmon survival in the wild. Here, we detected a genetic correlation whereby survival until the second spawning event was linked to the genome region that is also associated with sea age at maturity (Ayllon *et al.*, 2015; Barson *et al.*, 2015), another important fitness-linked life history trait. The allele associated with a higher prevalence of repeat spawning was also linked to earlier first-time maturation (i.e., the *E* allele in Barson *et al.,* 2015). This genetic correlation suggests that individuals with the *E* allele increase their fitness (relative to the *L* allele, associated with late maturation) not only by means of spending less time at sea, hence lowering the risk of mortality and shorter generation time, but also by increasing reproductive success by participating in multiple spawning events.

The genetic correlation between an iteroparous reproductive strategy and earlier first-time spawning is in accordance with energy allocation similarities between these two life history strategies as predicted by life history theory: investment in reproduction is higher in semelparous organisms/individuals (a single reproduction event before death) compared to those with and an iteroparous reproductive strategy (repeat spawning in this case, Crespi & Teo, 2002). Reproductive investment is indeed higher in large, late-maturing individuals who have spent two or more years at sea prior to first spawning, as quantified by a higher proportion of energy deposited to gonadal organs (Jonsson *et al.*, 1997; Jonsson & Jonsson, 2003), while gonadal investment in repeat spawners is lower (Niemelä *et al.*, 2000; Fleming & Einum, 2010). Furthermore, the *vgll3* gene, the most likely gene candidate in the region associated with both traits, regulates adipose content and body weight in mice (Halperin *et al.*, 2013), both of which are postulated to regulate maturation age and repeat spawning in Atlantic salmon (Friedland & Haas, 1996; Jonsson *et al.*, 1997; Taranger *et al.*, 2010). Our results support the notion that the genetic variation in these two life history strategies may be mediated by similar a physiological cascade controlled, at least in part, by the same genomic region.

Even though sea age at maturity and repeat spawning are genetically correlated and regulated by similar physiological cascades, it is unclear if the same SNP is linked to trait variation in both traits (i.e., pleiotropy in the strict sense, see Wagner and Zhang, 2011) or if the association is a result of a causal SNPs being closely linked in the region. Although *vgll3*_*Met54Thr*_, a SNP causing a potentially functionally important missense substitution in the *vgll3* gene (Barson *et al.*, 2015), appears to be a prime candidate for a causal association to iteroparity, other candidate SNPs in the region were not conclusively ruled out as the region of highest association. Likewise, causality between SNPs in the region and age at maturity is yet to be tested further (Barson *et al.*, 2015). Narrowing down the causal SNP will be important to gain a better understanding of the physiological mechanisms underlying the trait variation and, in this particular case, the nature of the genetic correlation between the two traits. This may be particularly important in estimating potential evolutionary trajectories of life history co-evolution in salmon (Etterson & Shaw, 2001; Steppan *et al.*, 2002; Conner *et al.*, 2011). For example, pleiotropic genetic correlations are harder to break up than linkages that are non-pleiotropic and physically distant from each other, e.g., if the genetic correlation is maladaptive (Mackay, 2001; Gardner & Latta, 2007; Mackay *et al.*, 2009). The nature of the linkage should be further investigated to better evaluate the co-evolutionary potential between life history traits over contemporary time scales (Gardner & Latta, 2007).

In most documented cases, the evolutionary potential of genetically correlated fitness traits is constrained by the opposing directional response of correlated trait values to the selection gradient (e.g., (Etterson & Shaw, 2001; Charmantier *et al.*, 2006; Theriault *et al.*, 2007; Fordyce & Nice, 2008; Nussey *et al.*, 2008; Simon *et al.*, 2016), which is predicted to help maintain variation in fitness traits (Roff, 1996). In our study, the trait variation within a population was not maintained by opposing selective directions of correlated trait values, since sea age at maturity is not under directional selection, but variation is being maintained by balancing selection between younger and older maturation ages (Barson *et al.*, 2015). Therefore, the dynamics underlying the genetic covariation cannot be explained by antagonistic trait correlation. Assuming that iteroparity in Atlantic salmon is linked to higher reproductive success, a genetic correlation between an iteroparous reproductive strategy and sea age at maturity would reinforce the prevalence of a younger age structure; hence, a younger population age structure would be predicted than if the genetic correlation were not accounted for. On the other hand, it is not clear if repeat spawning is always linked to higher fitness. In steelhead salmon (*Oncorhynchus mykiss*), iteroparity is linked to increased overall reproductive success, but the first-time reproductive output of repeat spawners is less than that of semelparous individuals at the same age (Christie *et al.*, 2018), suggesting that a fitness advantage may be reversed if conditions promoting post-spawning survival at sea deteriorate (e.g., Chaput & Benoit, 2012).

There was a contradiction between the sex-specific phenotypic patterns of repeat spawning compared to the observed genetic basis of repeat spawning variation (Table 1). Females tend to have a higher level of iteroparity than males (Niemelä *et al.*, 2006a), but this phenotypic difference was not translated into a genetic association at the *vgll3*_*Met54Thr*_ locus, which is independent of sex. This lends support to the notion that the sex-specific differences in repeat spawning may be attributed to sex-specific behavioural differences rather than physiological differences (e.g., Niemelä *et al.*, 2006a). For example, despite the much lower gonadosomatic index of male gonads (Jonsson *et al.*, 1991b; 1997), males are more aggressive and more likely to attempt reproduction until exhausting themselves to the point of death and/or sustain more damage during reproduction (Jonsson *et al.*, 1997), while female reproductive effort is limited to egg deposition. From this perspective, our results are consistent with a notion that genetic variation in the *vgll3*_*Met54Thr*_ locus is associated with physiological mechanisms that are not linked to sex-specific differences in behaviour. On the other hand, the genetic regulation in repeat spawning prevalence was substantially stronger in later-maturing fish. Stronger physiological constraints dramatically impede survival in older sea age groups, likely as a result of higher energetic loss during maturation and spawning, as well as relatively higher maintenance metabolism (Jonsson *et al.*, 1991a; Jonsson *et al.*, 1997; Metcalfe *et al.*, 2015). The genetic correlation between maturation age and repeat spawning prevalence suggests the same energetic constraints underlying the physiological basis of both traits may be controlled by the *vgll3* locus. Overall, understanding the interplay between the phenotypic and genetic correlation between repeat spawning and other phenotypic indices (sex and sea age at maturity) will provide further insights into the functional basis of the association between these traits.

Post-smolt growth at sea is an important determinant of survival in salmon (e.g., Friedland *et al.,* 2005) and has been shown to be positively correlated with early age at maturity (Friedland & Haas, 1996). Likewise, better growth conditions at sea are correlated to post-spawning survival (Chaput & Benoit, 2012). Here, unlike the first-time spawners, we showed that size did not differ between repeat spawners with different *vgll3* genotypes (Table 2). However, it is unclear from the terminal length data at which life history stage the genotype-specific size differences observed in first-time spawners were offset in repeat spawners. Two plausible explanations exist. i) Surviving the first spawning event is length-dependent for individuals with the *EE* genotype but not for those with the *LL* genotype. ii) Fish with the *EE* genotype grow faster than *LL* individuals after the first spawning event. By measuring the length of surviving post-spawned fish in the Teno River prior to their second sea migration, Niemelä *et al.* (2000) showed that repeat spawners were, on average, larger than first-time spawners. This information is consistent with the first possibility, that larger fish with the *EE* genotype are likely to survive the first spawning, while this survival is length independent for fish with the *LL* genotype.

To investigate the second possibility further, i.e., if post-spawning growth in *EE* individuals is higher than in *LL* individuals, we performed a *post hoc* analysis and quantified post-spawning scale growth as a proxy for growth, using a similar modelling framework as for other analyses (Appendix S1, Figure S3). This analysis appeared to be underpowered, as many models, including null structures, exhibited similarly high explanatory powers (Δ*AICc*<2 in the most parsimonious models) (see Table S9). However, all parsimonious models rejected the scenario of higher post-spawning growth being linked to the *E* allele (Table S9), refuting that the better post-spawn growth in the *EE* genotype explains growth patterns in repeat spawners. Finally, in all plausible models, post-spawning growth was negatively correlated to sea age (Table S9), further suggesting that older sea age groups had very limited post-spawning growth. In fact, when post- and pre-spawning scale growth was plotted over total length separately, a correlation between growth after first spawning and total length was only observed in 1S1 fish. This further illustrates the dependence of growth after first spawning on sea age (Figure S4, see also Supplementary Information).

Overall, this *ad hoc* analysis confirmed that the genotype-dependent differences observed between repeat spawners and first-time spawners arose prior to first migration, rather than after. In addition, we further showed growth during the post-spawning period was limited in later-maturing fish. This was probably as a result of the higher energy demands of larger, older maturing fish to restore and maintain their body mass after a reproduction event (Jonsson *et al.*, 1997). Individuals maturing later invest more energy in reproduction (Jonsson *et al.*, 1997), and therefore, it might be more difficult to maintain positive energy balance after the first reproduction event. Relatively little is known on the dynamics of iteroparity in Atlantic salmon (Thorstad *et al.*, 2010), and our results thus provide valuable insight on the growth dynamics of alternative reproductive tactics.

The abundance of Atlantic salmon populations has declined substantially over the past 40 years and is at an all-time low level (ICES; Chaput, 2012). This decline is coupled with demographic changes, mostly towards younger age structure (Chaput, 2012), and global climate change will likely affect population structure further (Friedland *et al.*, 2005; Friedland *et al.*, 2009; Hedger *et al.*, 2013; Mills *et al.*, 2013; Piou & Prevost, 2013; Jonsson *et al.*, 2016). Many salmon management regimes consider the repeat spawner phenotype a suitable substitute for dwindling numbers of large, late-maturing, first-time spawning individuals, and their presence in populations has been linked with improved genetic stability (Hatch *et al.*, 2004; Niemelä *et al.*, 2006a; Narum *et al.*, 2008; Seamons & Quinn, 2009; Chaput & Benoit, 2012; Reid & Chaput, 2012). However, the genetic evidence presented here suggests that increases in repeat spawner numbers may be associated with decreasing age structure of first-time spawners, and directional selection changing sea age composition (e.g., in response to environmental changes) may result in changes in repeat spawner composition, which may influence the demographic structure further. Despite the fact that genetic effects have been increasingly included in demographic models (i.e., demo-genetic models, i.e., (Dunlop *et al.*, 2009; Reed *et al.*, 2011; Piou & Prévost, 2012), accurately accounting for genetic architecture, e.g., (Kuparinen & Hutchings, 2017), gene-trait association, genetic correlations, and functional links between correlated traits would provide more realistic predictions, which could improve future conservation and management efforts.

## Acknowledgements

We acknowledge the fishers of the Teno River who contributed scales and phenotypic information to the Natural Resources Institute Finland (LUKE). Scale analyses were carried out by Jari Haantie. Jorma Kuusela is thanked for help with scale selection, analysis. Meri Lindqvist, Heli Junes, Jan Gerwin, Minna Kuusela and Kristiina Haapanen are thanked for laboratory assistance. This study was supported by grants from the Maj and Tor Nessling Foundation (project number 201600445 to TA), and Finnish Academy (grant number 318939 to TA, 286334 to JE and, 284941, 307593, 302873 to CRP). Victoria L. Pritchard is thanked for analysis suggestions. Data and codes related to model optimization and parameters are available in the Dryad Digital Repository: XXX-XXX.

## References

Agrawal, A.F. & Stinchcombe, J.R. 2009. How much do genetic covariances alter the rate of adaptation? Proc. Biol. Sci. 276: 1183–1191.

Aykanat, T., Johnston, S.E., Orell, P., Niemela, E., Erkinaro, J. & Primmer, C.R. 2015. Low but significant genetic differentiation underlies biologically meaningful phenotypic divergence in a large Atlantic salmon population. Mol. Ecol. 24: 5158–5174.

Aykanat, T., Lindqvist, M., Pritchard, V.L. & Primmer, C.R. 2016. From population genomics to conservation and management: a workflow for targeted analysis of markers identified using genome-wide approaches in Atlantic salmon Salmo salar. J. Fish. Biol. 89:2658–2679.

Ayllon, F., Kjaerner-Semb, E., Furmanek, T., Wennevik, V., Solberg, M.F., Dahle, G., Taranger, G.L., Glover, K.A., Almen, M.S., Rubin, C.J., Edvardsen, R.B. & Wargelius, A. 2015. The vgll3 Locus Controls Age at Maturity in Wild and Domesticated Atlantic Salmon (Salmo salar L.) Males. PLoS Genet. 11: e1005628.

Barson, N.J., Aykanat, T., Hindar, K., Baranski, M., Bolstad, G.H., Fiske, P., Jacq, C., Jensen, A.J., Johnston, S.E., Karlsson, S., Kent, M., Moen, T., Niemela, E., Nome, T., Naesje, T.F., Orell, P., Romakkaniemi, A., Saegrov, H., Urdal, K., Erkinaro, J., Lien, S. & Primmer, C.R. 2015. Sex-dependent dominance at a single locus maintains variation in age at maturity in salmon. Nature 528: 405–408.

Berenos, C., Ellis, P.A., Pilkington, J.G. & Pemberton, J.M. 2014. Estimating quantitative genetic parameters in wild populations: a comparison of pedigree and genomic approaches. Mol. Ecol. 23: 3434–3451.

Bulik-Sullivan, B., Finucane, H.K., Anttila, V., Gusev, A., Day, F.R., Loh, P.R., ReproGen, C., Psychiatric Genomics, C., Genetic Consortium for Anorexia Nervosa of the Wellcome Trust Case Control, C., Duncan, L., Perry, J.R., Patterson, N., Robinson, E.B., Daly, M.J., Price, A.L. & Neale, B.M. 2015. An atlas of genetic correlations across human diseases and traits. Nat. Genet. 47: 1236–1241.

Campbell, N.R., Harmon, S.A. & Narum, S.R. 2015. Genotyping-in-Thousands by sequencing (GT-seq): A cost effective SNP genotyping method based on custom amplicon sequencing. Mol. Ecol. Resour. 15: 855-867.

Careau, V., Reale, D., Humphries, M.M. & Thomas, D.W. 2010. The pace of life under artificial selection: personality, energy expenditure, and longevity are correlated in domestic dogs. Am. Nat. 175:753-758.

Carlson, S.M. & Seamons, T.R. 2008. A review of quantitative genetic components of fitness in salmonids: implications for adaptation to future change. Evol. Appl. 1:222-238.

Chaput, G. 2012. Overview of the status of Atlantic salmon (Salmo salar) in the North Atlantic and trends in marine mortality. Ices J. Mar. Sci. 69: 1538–1548.

Chaput, G. & Benoit, H.P. 2012. Evidence for bottom-up trophic effects on return rates to a second spawning for Atlantic salmon (Salmo salar) from the Miramichi River, Canada. Ices J. Mar. Sci. 69: 1656-1667.

Charmantier, A., Perrins, C., McCleery, R.H. & Sheldon, B.C. 2006. Quantitative genetics of age at reproduction in wild swans: Support for antagonistic pleiotropy models of senescence. Proc. Natl. Acad. Sci. USA 103: 6587-6592.

Chen, C.Y., Misztal, I., Aguilar, I., Tsuruta, S., Meuwissen, T.H., Aggrey, S.E., Wing, T. & Muir, W.M. 2011. Genome-wide marker-assisted selection combining all pedigree phenotypic information with genotypic data in one step: An example using broiler chickens. J. Anim. Sci. 89 23-28.

Chevin, L.M., Martin, G. & Lenormand, T. 2010. Fisher’s model and the genomics of adaptation: restricted pleiotropy, heterogenous mutation, and parallel evolution. Evolution 64: 3213–3231.

Chirgwin, E., Monro, K., Sgro, C.M. & Marshall, D.J. 2015. Revealing hidden evolutionary capacity to cope with global change. Glob. Chang. Biol. 21: 3356–3366.

Christie, M.R., McNickle, G.G., French, R.A. & Blouin, M.S. 2018. Life history variation is maintained by fitness trade-offs and negative frequency-dependent selection. Proc Natl Acad Sci U S A.

Clements, M.N., Clutton-Brock, T.H., Guinness, F.E., Pemberton, J.M. & Kruuk, L.E. 2011. Variances and covariances of phenological traits in a wild mammal population. Evolution 65: 788–801.

Conner, J.K., Karoly, K., Stewart, C., Koelling, V.A., Sahli, H.F. & Shaw, F.H. 2011. Rapid independent trait evolution despite a strong pleiotropic genetic correlation. Am Nat 178: 429–441.

Crespi, B.J. & Teo, R. 2002. Comparative phylogenetic analysis of the evolution of semelparity and life history in salmonid fishes. Evolution 56: 1008–1020.

Dunlop, E.S., Heino, M. & Dieckmann, U. 2009. Eco-genetic modeling of contemporary life-history evolution. Ecol. Appl. 19: 1815–1834.

Duputie, A., Massol, F., Chuine, I., Kirkpatrick, M. & Ronce, O. 2012. How do genetic correlations affect species range shifts in a changing environment? Ecol. Lett. 15: 251–259.

Erkinaro, J., Niemelä, E., Vähä, J.-P., Primmer, C.R., Brørs, S. & Hassinen, E. 2010. Distribution and biological characteristics of escaped farmed salmon in a major subarctic wild salmon river: implications for monitoring. Can. J. Fish. Aquat. Sci. 67: 130-142.

Erkinaro, J., Czorlich, Y., Orell, P., Kuusela, J., Falkegård, M., Länsman, M., Pulkkinen, H., Primmer, C.R. & Niemelä, E. 2018. Life history variation across four decades in a diverse population complex of Atlantic salmon in a large subarctic river. Can. J. Fish. Aquat. Sci..

Etterson, J.R. & Shaw, R.G. 2001. Constraint to adaptive evolution in response to global warming. Science294: 151–154.

Fleming, I.A. 1996. Reproductive strategies of Atlantic salmon: ecology and evolution. Rev. Fish. Biol. Fisher. 6: 379-416.

Fleming, I.A. & Einum, S. 2010. Reproductive Ecology: A Tale of Two Sexes. In: Atlantic Salmon Ecology, pp. 33–65. Wiley-Blackwell.

Fordyce, J.A. & Nice, C.C. 2008. Antagonistic, stage-specific selection on defensive chemical sequestration in a toxic butterfly. Evolution 62: 1610–1617.

Friedland, K.D. & Haas, R.E. 1996. Marine post-smolt growth and age at maturity of Atlantic salmon. J. Fish. Biol. 48: 1-15.

Friedland, K.D., Chaput, G. & MacLean, J.C. 2005. The emerging role of climate in post-smolt growth of Atlantic salmon. Ices J. Mar. Sci. 62: 1338–1349.

Friedland, K.D., MacLean, J.C., Hansen, L.P., Peyronnet, A.J., Karlsson, L., Reddin, D.G., Maoileidigh, N.O. & McCarthy, J.L. 2009. The recruitment of Atlantic salmon in Europe. Ices J. Mar. Sci. 66: 289–304.

Gardner, K.M. & Latta, R.G. 2007. Shared quantitative trait loci underlying the genetic correlation between continuous traits. Mol. Ecol. 16: 4195–4209.

Gratten, J. & Visscher, P.M. 2016. Genetic pleiotropy in complex traits and diseases: implications for genomic medicine. Genome Med. 8: 78.

Griswold, C.K. & Whitlock, M.C. 2003. The genetics of adaptation: the roles of pleiotropy, stabilizing selection and drift in shaping the distribution of bidirectional fixed mutational effects. Genetics 165: 2181–2192.

Halperin, D.S., Pan, C., Lusis, A.J. & Tontonoz, P. 2013. Vestigial-like 3 is an inhibitor of adipocyte differentiation. J. Lipid. Res. 54: 473–481.

Hammerschlag, A.R., Stringer, S., de Leeuw, C.A., Sniekers, S., Taskesen, E., Watanabe, K., Blanken, T.F., Dekker, K., Te Lindert, B.H.W., Wassing, R., Jonsdottir, I., Thorleifsson, G., Stefansson, H., Gislason, T., Berger, K., Schormair, B., Wellmann, J., Winkelmann, J., Stefansson, K., Oexle, K., Van Someren, E.J.W. & Posthuma, D. 2017. Genome-wide association analysis of insomnia complaints identifies risk genes and genetic overlap with psychiatric and metabolic traits. Nat. Genet. 49: 1584–1592.

Hatch, D.R., Branstetter, R.D., Whiteaker, J., Blodgett, J., Bosch, B., Fast, D. & Newsome, T. 2004. Kelt Reconditioning: A Research Project to Enhance Iteroparity in Columbia Basin Steelhead (Oncorhynchus mykiss). Annual Report to U.S. Department of Energy, Bonneville Power Administration, Project No. 2000–017. Portland, OR: Bonneville Power Administration.

Hedger, R.D., Naesje, T.F., Fiske, P., Ugedal, O., Finstad, A.G. & Thorstad, E.B. 2013. Ice-dependent winter survival of juvenile Atlantic salmon. Ecol. Evol. 3: 523–535.

Hellmann, J.J. & Pineda-Krch, M. 2007. Constraints and reinforcement on adaptation under climate change: Selection of genetically correlated traits. Biological Conservation 137: 599–609.

Houle, D. 1991. Genetic Covariance of Fitness Correlates - What Genetic Correlations Are Made of and Why It Matters. Evolution 45: 630–648.

ICES. 2011. Report of the Workshop on Age Determination of Salmon (WKADS), 18L20 January 2011, Galway, Ireland. ICES CM 2011/ACOM:44. |p67 pp.

ICES. 2018. Report of the Working Group on North Atlantic Salmon (WGNAS), 4–13 April 2018, Woods Hole, MA, USA. ICES CM 2018/ACOM:21. 386 pp.

Johnston, S.E., Orell, P., Pritchard, V.L., Kent, M.P., Lien, S., Niemela, E., Erkinaro, J. & Primmer, C.R. 2014. Genome-wide SNP analysis reveals a genetic basis for sea-age variation in a wild population of Atlantic salmon (Salmo salar). Mol. Ecol. 23: 3452–3468.

Jonsson, B., Jonsson, N. & Albretsen, J. 2016. Environmental change influences the life history of salmon Salmo salar in the North Atlantic Ocean. J Fish Biol.

Jonsson, N., Hansen, L.P. & Jonsson, B. 1991a. Variation in Age, Size and Repeat Spawning of Adult Atlantic Salmon in Relation to River Discharge. J. Anim. Ecol. 60: 937–947.

Jonsson, N., Jonsson, B. & Hansen, L.P. 1991b. Energetic Cost of Spawning in Male and Female Atlantic Salmon (Salmo salar L). J. Fish. Biol. 39: 739–744.

Jonsson, N., Jonsson, B. & Hansen, L.P. 1997. Changes in proximate composition and estimates of energetic costs during upstream migration and spawning in Atlantic salmon Salmo salar. J. Anim. Ecol. 66: 425–436.

Jonsson, N. & Jonsson, B. 2003. Energy allocation among developmental stages, age groups, and types of Atlantic salmon (Salmo salar) spawners. Can. J. Fish. Aquat. Sci. 60: 506–516.

Kadarmideen, H.N., Thompson, R., Coffey, M.P. & Kossaibati, M.A. 2003. Genetic parameters and evaluations from single- and multiple-trait analysis of dairy cow fertility and milk production. Livestock Production Science 81: 183–195.

Kause, A., Quinton, C., Airaksinen, S., Ruohonen, K. & Koskela, J. 2011. Quality and production trait genetics of farmed European whitefish, Coregonus lavaretus. J Anim Sci 89: 959–971.

Kruuk, L.E.B., Clutton-Brock, T.H., Slate, J., Pemberton, J.M., Brotherstone, S. & Guinness, F.E. 2000. Heritability of fitness in a wild mammal population. Proc. Natl. Acad. Sci. USA 97: 698–703.

Kuparinen, A. & Hutchings, J.A. 2017. Genetic architecture of age at maturity can generate divergent and disruptive harvest-induced evolution. Philos. Trans. R. Soc. Lond. B Biol. Sci. 372.

Lande, R. 1979. Quantitative Genetic Analysis of Multivariate Evolution, Applied to Brain:Body Size Allometry. Evolution 33: 402–416.

Lande, R. 1982. A Quantitative Genetic Theory of Life-History Evolution. Ecology 63: 607–615.

Lande, R. & Arnold, S.J. 1983. The Measurement of Selection on Correlated Characters. Evolution 37: 1210–1226.

Lane, J.E., Kruuk, L.E., Charmantier, A., Murie, J.O., Coltman, D.W., Buoro, M., Raveh, S. & Dobson, F.S. 2011. A quantitative genetic analysis of hibernation emergence date in a wild population of Columbian ground squirrels. J Evol Biol 24: 1949–1959.

Lee, S.H., Yang, J., Goddard, M.E., Visscher, P.M. & Wray, N.R. 2012. Estimation of pleiotropy between complex diseases using single-nucleotide polymorphism-derived genomic relationships and restricted maximum likelihood. Bioinformatics 28: 2540–2542.

Mackay, T.F., Stone, E.A. & Ayroles, J.F. 2009. The genetics of quantitative traits: challenges and prospects. Nat. Rev. Genet. 10: 565–577.

Mackay, T.F.C. 2001. The genetic architecture of quantitative traits. Annu. Rev. Genet. 35: 303–339.

Metcalfe, N.B., Van Leeuwen, T.E. & Killen, S.S. 2016. Does individual variation in metabolic phenotype predict fish behaviour and performance? J. Fish. Biol. 88:298–321.

Mills, K.E., Pershing, A.J., Sheehan, T.F. & Mountain, D. 2013. Climate and ecosystem linkages explain widespread declines in North American Atlantic salmon populations. Glob. Chang. Biol. 19: 3046–3061.

Narum, S.R., Hatch, D., Talbot, A.J., Moran, P. & Powell, M.S. 2008. Iteroparity in complex mating systems of steelhead Oncorhynchus mykiss (Walbaum). J. Fish. Biol. 72: 45–60.

Niemelä, E., Makinen, T.S., Moen, K., Hassinen, E., Erkinaro, J., Lansman, M. & Julkunen, M. 2000. Age, sex ratio and timing of the catch of kelts and ascending Atlantic salmon in the subarctic River Teno. J. Fish. Biol. 56: 974–985.

Niemelä, E., Erkinaro, J., Julkunen, M., Hassinen, E., Lansman, M. & Brors, S. 2006a. Temporal variation in abundance, return rate and life histories of previously spawned Atlantic salmon in a large subarctic river. J. Fish. Biol. 68: 1222–1240.

Niemelä, E., Orell, P., Erkinaro, J., Dempson, J.B., BrØrs, S., Svenning, M.A. & Hassinen, E. 2006b. Previously spawned Atlantic salmon ascend a large subarctic river earlier than their maiden counterparts. J. Fish. Biol. 69: 1151–1163.

Nussey, D.H., Wilson, A.J., Morris, A., Pemberton, J., Clutton-Brock, T. & Kruuk, L.E. 2008. Testing for genetic trade-offs between early-and late-life reproduction in a wild red deer population. Proc Biol Sci 275: 745–750.

Orr, H.A. 2000. Adaptation and the cost of complexity. Evolution 54: 13–20.

Ozerov, M., Vasemagi, A., Wennevik, V., Niemela, E., Prusov, S., Kent, M. & Vaha, J.P. 2013. Cost-effective genome-wide estimation of allele frequencies from pooled DNA in Atlantic salmon (Salmo salar L.). BMC Genomics 14: 12.

Penney, Z.L. & Moffitt, C.M. 2013. Histological assessment of organs in sexually mature and post-spawning steelhead trout and insights into iteroparity. Rev. Fish Biol. Fisher. 24: 781–801.

Piou, C. & Prévost, E. 2012. A demo-genetic individual-based model for Atlantic salmon populations: Model structure, parameterization and sensitivity. Ecol. Model. 231: 37–52.

Piou, C. & Prevost, E. 2013. Contrasting effects of climate change in continental vs. oceanic environments on population persistence and microevolution of Atlantic salmon. Glob. Chang. Biol. 19: 711–723.

Pritchard, V.L., Makinen, H., Vaha, J.P., Erkinaro, J., Orell, P. & Primmer, C.R. 2018. Genomic signatures of fine-scale local selection in atlantic salmon suggest involvement of sexual maturation, energy homeostasis, and immune defence-related genes. Mol. Ecol.

Reed, T.E., Schindler, D.E., Hague, M.J., Patterson, D.A., Meir, E., Waples, R.S. & Hinch, S.G. 2011. Time to evolve? Potential evolutionary responses of fraser river sockeye salmon to climate change and effects on persistence. PLoS One 6: e20380.

Reid, J.E. & Chaput, G. 2012. Spawning history influence on fecundity, egg size, and egg survival of Atlantic salmon (Salmo salar) from the Miramichi River, New Brunswick, Canada. Ices J. Mar. Sci. 69: 1678–1685.

Ritland, K. 1996. A Marker-based method for inferences about quantitative inheritance in natural populations. Evolution 50: 1062–1073.

Robinson, M.R., Wilson, A.J., Pilkington, J.G., Clutton-Brock, T.H., Pemberton, J.M. & Kruuk, L.E. 2009. The impact of environmental heterogeneity on genetic architecture in a wild population of Soay sheep. Genetics 181: 1639–1648.

Robinson, M.R., Santure, A.W., Decauwer, I., Sheldon, B.C. & Slate, J. 2013. Partitioning of genetic variation across the genome using multimarker methods in a wild bird population. Mol. Ecol. 22: 3963–3980.

Roff, D.A. 1996. The Evolution of Genetic Correlations: An Analysis of Patterns. Evolution 50: 1392–1403.

Sanchez-Guillen, R.A., Wellenreuther, M. & Cordero Rivera, A. 2012. Strong asymmetry in the relative strengths of prezygotic and postzygotic barriers between two damselfly sister species. Evolution 66: 690–707.

Santure, A.W., De Cauwer, I., Robinson, M.R., Poissant, J., Sheldon, B.C. & Slate, J. 2013. Genomic dissection of variation in clutch size and egg mass in a wild great tit (Parus major) population. Mol. Ecol. 22: 3949–3962.

Schnurr, T.M., Gjesing, A.P., Sandholt, C.H., Jonsson, A., Mahendran, Y., Have, C.T., Ekstrom, C.T., Bjerregaard, A.L., Brage, S., Witte, D.R., Jorgensen, M.E., Aadahl, M., Thuesen, B.H., Linneberg, A., Eiberg, H., Pedersen, O., Grarup, N., Kilpelainen, T.O. & Hansen, T. 2016. Genetic Correlation between Body Fat Percentage and Cardiorespiratory Fitness Suggests Common Genetic Etiology. PLoS One 11: e0166738.

Seamons, T.R. & Quinn, T.P. 2009. Sex-specific patterns of lifetime reproductive success in single and repeat breeding steelhead trout (Oncorhynchus mykiss). Behav. Ecol. Sociobiol. 64: 505–513.

Sheldon, B.C., Kruuk, L.E. & Merila, J. 2003. Natural selection and inheritance of breeding time and clutch size in the collared flycatcher. Evolution 57: 406–420.

Simon, M.N., Machado, F.A. & Marroig, G. 2016. High evolutionary constraints limited adaptive responses to past climate changes in toad skulls. Proc. Biol. Sci. 283.

Stearns, S., Dejong, G. & Newman, B. 1991. The Effects of Phenotypic Plasticity on Genetic Correlations. Trends Ecol. Evol. 6: 122–126.

Steppan, S.J., Phillips, P.C. & Houle, D. 2002. Comparative quantitative genetics: evolution of the G matrix. Trends Ecol. Evol. 17: 320–327.

Storz, J.F., Bridgham, J.T., Kelly, S.A. & Garland, T., Jr. 2015. Genetic approaches in comparative and evolutionary physiology. Am. J. Physiol. Regul. Integr. Comp. Physiol. 309: R197–214.

Taranger, G.L., Carrillo, M., Schulz, R.W., Fontaine, P., Zanuy, S., Felip, A., Weltzien, F.A., Dufour, S., Karlsen, O., Norberg, B., Andersson, E. & Hansen, T. 2010. Control of puberty in farmed fish. Gen. Comp. Endocrinol. 165: 483–515.

Theriault, V., Garant, D., Bernatchez, L. & Dodson, J.J. 2007. Heritability of life-history tactics and genetic correlation with body size in a natural population of brook charr (Salvelinus fontinalis). J. Evol. Biol. 20: 2266–2277.

Thorstad, E.B., Whoriskey, F., Rikardsen, A.H. & Aarestrup, K. 2010. Aquatic Nomads: The Life and Migrations of the Atlantic Salmon. In: Atlantic Salmon Ecology, pp. 1–32. Wiley-Blackwell.

Vähä, J.-P., Erkinaro, J., Falkegård, M., Orell, P. & Niemelä, E. 2017. Genetic stock identification of Atlantic salmon and its evaluation in a large population complex. Can. J. Fish. Aquat. Sci. 74: 327–338.

Vähä, J.P., Erkinaro, J., Niemelä, E. & Primmer, C.R. 2007. Life-history and habitat features influence the within-river genetic structure of Atlantic salmon. Mol. Ecol. 16: 2638–2654.

Wagner, G.P. 1989. Multivariate mutation-selection balance with constrained pleiotropic effects. Genetics 122: 223–234.

Wagner, G.P. & Zhang, J. 2011. The pleiotropic structure of the genotype-phenotype map: the evolvability of complex organisms. Nat. Rev. Genet. 12: 204–213.

Walters, C. & Maguire, J.J. 1996. Lessons for stock assessment from the northern cod collapse. Rev. Fish Biol. Fisher. 6: 125–137.

Wang, Z., Liao, B.Y. & Zhang, J. 2010. Genomic patterns of pleiotropy and the evolution of complexity. Proc. Natl. Acad. Sci. USA 107: 18034–18039.

Weigel, K.A., VanRaden, P.M., Norman, H.D. & Grosu, H. 2017. A 100-Year Review: Methods and impact of genetic selection in dairy cattle-From daughter-dam comparisons to deep learning algorithms. J. Dairy. Sci. 100: 10234–10250.

Wilson, A.J., Reale, D., Clements, M.N., Morrissey, M.M., Postma, E., Walling, C.A., Kruuk, L.E. & Nussey, D.H. 2010. An ecologist’s guide to the animal model. J. Anim. Ecol. 79: 13–26.

Yang, J., Benyamin, B., McEvoy, B.P., Gordon, S., Henders, A.K., Nyholt, D.R., Madden, P.A., Heath, A.C., Martin, N.G., Montgomery, G.W., Goddard, M.E. & Visscher, P.M. 2010. Common SNPs explain a large proportion of the heritability for human height. Nat. Genet. 42: 565–569.

